# Purified cytosolic crystals from *Bacillus thuringiensis* as a novel active pharmaceutical ingredient (API)

**DOI:** 10.1101/2022.05.20.492900

**Authors:** Jeffrey Chicca, Nicholas R. Cazeault, Florentina Rus, Ambily Abraham, Carli Garceau, Hanchen Li, Samar M. Atwa, Kelly Flanagan, Ernesto R. Soto, Mary S. Morrison, David Gazzola, Yan Hu, David R. Liu, Martin K. Nielsen, Joseph F. Urban, Gary R. Ostroff, Raffi V. Aroian

## Abstract

*Bacillus thuringiensis* or Bt is a Gram-positive soil bacterium, widely and safely applied in the environment as an insecticide for combatting insect pests that damage crops and vector diseases. Dominant active ingredients made by Bt are insect-killing crystal (Cry) proteins released as crystalline inclusions upon bacterial sporulation. Some Bt Cry proteins, *e*.*g*., Cry5B, target nematodes (roundworms) and show exceptional promise as anthelmintics (cures for parasitic nematode diseases). We have recently described IBaCC (for **I**nactivated **Ba**cteria with **C**ytosolic **C**rystal(s)) in which bioactive Bt Cry crystals (containing Cry5B) are fully contained within the cytosol of dead bacterial ghosts. Here we demonstrate that these IBaCC-trapped Cry5B crystals can be liberated and purified away from cellular constituents yielding **P**urified **C**ytosolic **C**rystals (PCC). Cry5B PCC contains ∼95% Cry5B protein out of the total protein content. Cry5B PCC is highly bioactive against parasitic nematode larvae and adults *in vitro*. Cry5B PCC is also highly active *in vivo* against experimental human hookworm and *Ascaris* infections in rodents. The process was scaled up to the 100 liter scale to produce PCC for a pilot study to treat two foals infected with the Ascarid, *Parascaris* spp. Single dose Cry5B PCC brought the fecal egg counts of both foals to zero. These studies describe the process for the scalable production of purified Bt crystals and define a new active pharmaceutical ingredient form of Bt Cry proteins.

**NON-TECHNICAL IMPORTANCE PARAGRAPH:** *Bacillus thuringiensis* crystal proteins are widely and safely used as insecticides. Recent studies show they also can cure gastrointestinal parasitic worm (nematode) infections when ingested. However, reproducible, scalable, and practical techniques for purifying these proteins have been lacking. Here, we address this severe limitation and present scalable and practical methods for large-scale purification of potently bioactive *B. thuringiensis* crystals and crystal proteins. The resultant product, called Purified Cytosolic Crystals (PCC), is highly compatible with ingestible drug delivery and formulation. Furthermore, there are growing applications in agriculture and insect control where access to large quantities of purified crystal proteins are desirable and where these methods will find great utility.

## INTRODUCTION

*Bacillus thuringiensis* or Bt is a naturally occurring Gram positive soil bacterium and globally the most successfully and commonly used biologically produced insecticide, comprising 75% of the biopesticide market and a major insecticide used in organic farming (1, 2). Bt strains have evolved that kill lepidopteran (caterpillars) and coleopteran (beetles) pests that are highly damaging to agriculture and dipteran (mosquitoes, blackflies) pests that are important for vectoring major global diseases like malaria, River Blindness, and Dengue Fever (3–6). The main (but not only) insecticidal ingredients that Bt makes are three-domain crystal (Cry) proteins, so named because these proteins form large macromolecular crystalline inclusion bodies during bacterial sporulation (7). These proteins are safe and non-toxic to vertebrates, showing No Effect Levels (NOELs) in the > 1000 mg/kg range (8, 9). This safety factor has played an important role in the adoption and expansion of use of Cry proteins, including as expressed proteins in transgenic crops, *e*.*g*., to produce insect-resistant corn and cotton varieties that are now planted in >108 million hectares worldwide (10). In addition to targeting insects, some Bt Cry proteins also target nematodes, including intestinal parasites of humans, dogs, horses, sheep, and pigs and parasites of plants (11–17). Because nematodes are important parasites of humans, animals and plants, and because of their excellent safety record and profile, there is significant interest in using Bt Cry proteins to control nematode parasites.

For environmental applications, Bt crystals have always been deployed along with the live spores that produce them (*i*.*e*., as live spore crystal lysates; (18, 19)). Although the bacterium Bt is considered safe, it is in the same family as *Bacillus cereus*, a known enteric pathogen, and there are concerns that live Bt sprays could be related to sporadic food poisoning outbreaks (20–24). As such, in some places there are limits in terms of acceptable residual levels of Bt spores on foods sold (23, 24). There are also increasing environmental concerns with using live spores, *e*.*g*., Bt spore accumulation in non-target beneficial insects (25). For these and other reasons, using live Bt spore crystal lysates would be challenging from a regulatory and therapeutic vantage point as ingested anthelmintics (*e*.*g*., oral cures for intestinal nematodes) (14). The use of live spores also results in environmental contamination, which alters the natural bacterial composition and with potential implications for resistance (14). For all these considerations, we developed a novel form of Cry proteins delivered inside of dead bacteria. This novel form is called IBaCC for **I**nactivated **Ba**cteria with **C**ytosolic **C**rystal(s) (14). In IBaCC, Bt crystals are formed during vegetative growth/stationary phase in sporulation-defective Bt. The Bt crystals are trapped in the cytosol of the non-sporulated *Bacillus* bacteria (14). Such Bacilli cells are easily killed by food grade essential oils, while at the same time nematode-active Bt crystals left inside their dead bacterial ghosts fully retain bioactivity (14).

In the course of working with IBaCC, we found that the dead bacterial ghosts treated with essential oil can become somewhat brittle, raising the possibility that a simplified protocol might be developed in which Bt crystals could be isolated to high purity apart from other bacterial components. Here we describe such a simplified methodology for the isolation of **P**urified **C**ytosolic **C**rystals or PCC. Cry5B PCC was tested for bioactivity *in vitro* and *in vivo* against parasitic nematodes. We also scaled up purification of PCC utilizing a 100 liter culture of Bt Cry5B IBaCC. Use of Cry5B PCC from this scaled-up run for treatment of parasites in large animals (foals) is also presented.

## RESULTS

### IBaCC cells can be used to produce Purified Cytosolic Crystals (PCC)

We hypothesized that IBaCC cells could be cracked open and phase partitioning could be used to purify crystals away from soluble and insoluble bacterial debris. After significant trial and error, we developed a protocol that allowed for purification of crystals away from bacteria (Figure 1A; Materials and Methods). We found that recombinant IBaCC cells (killed with essential oil) expressing Cry5B crystals were relatively brittle and efficiently lysed by the combination of lysozyme and homogenization. Removal of cellular debris could be achieved by phase partitioning over hexane. If subsequent solubilization of the crystal protein from the crystal in solution was desired, an optional DNAse treatment step could be included to remove DNA fragments, often found in Bt crystals (26), which tend to keep the proteinaceous crystal intact in solution via ionic interactions.

**Figure 1.**
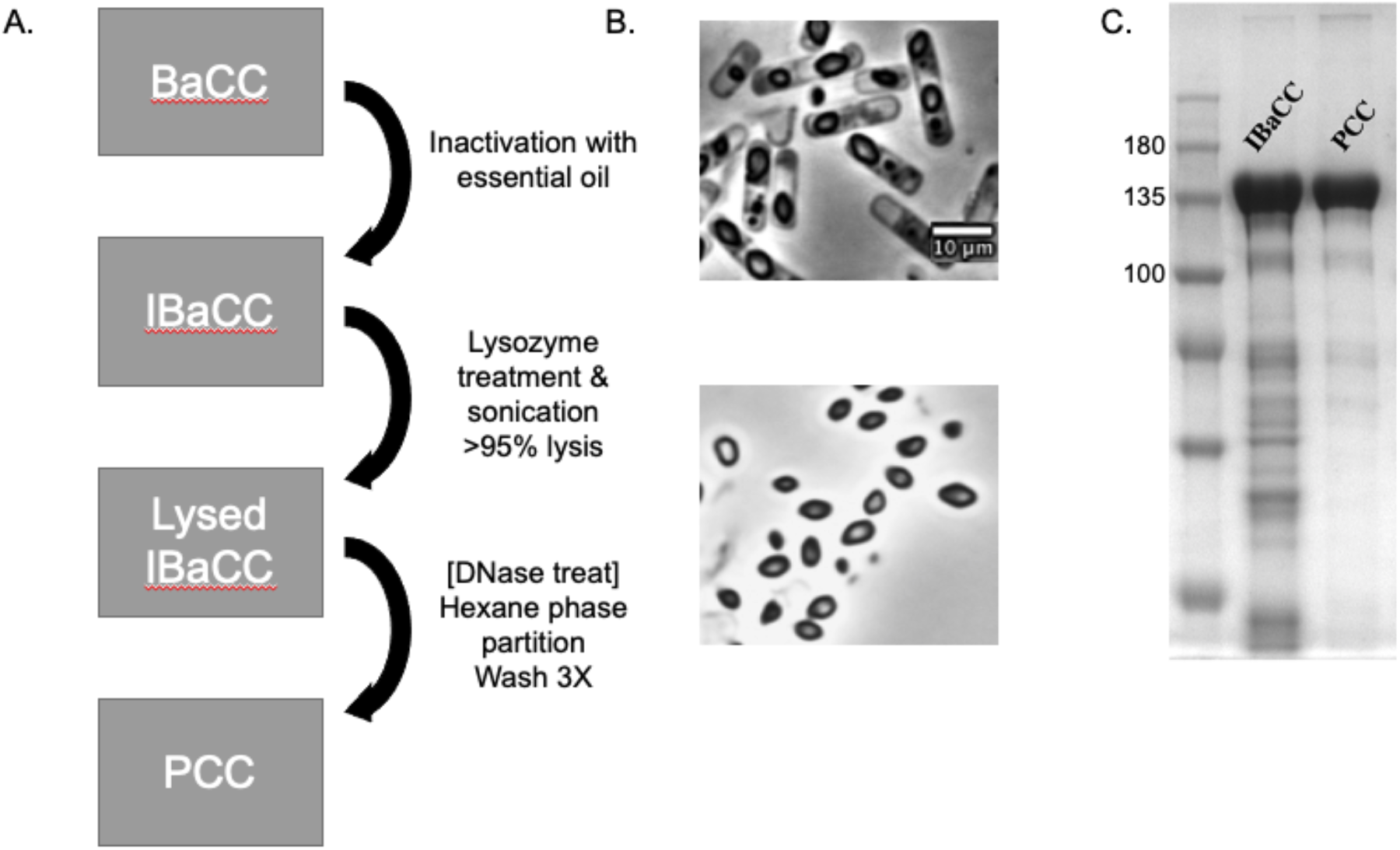
Purified Cytosolic Crystals (PCC). (A) A flow diagram of the process of turning BaCC into PCC. DNAse treatment is optional. (B) A comparison of IBaCC (upper panel) and PCC (lower panel) as visualized by using an Olympus BX60 microscope with 100x oil objective. The scale is the same in both panels. (C) SDS-PAGE visualization of Cry5B IBaCC and PCC. The left most lane includes protein standards (kDa).

The result of these process steps gives rise to Purified Cytosolic Crystals (PCC) as viewed by light microscopy (Figure 1B). By sodium dodecyl-sulfate polyacrylamide gel electrophoresis (SDS PAGE), a significant reduction in background protein gels can be seen (Figure 1C), with Cry5B making up 31% of Cry5B PCC by dry weight and 96% of the total protein (see Materials and Methods). We were also able to make Cry1Ac PCC from insecticidal Cry1Ac protein (see below).

### Cry5B PCC is bioactive *in vitro*

Blood feeding hookworms are amongst the most important of all human parasitic infections (27, 28). Roughly half a billion people have hookworm infestation, leading to an estimated disease burden of 4 million disability-adjusted life-years (DALYs) and economic losses of US $139 billion/year from lost work productivity. Human hookworms fall into two nematode genera, *Necator* and *Ancylostoma*, both of which are sensitive to Cry5B IBaCC (14).

We compared the bioactivity of Cry5B PCC and parent IBaCC against both genera of hookworm larvae using a larval development assay (29). Cry5B PCC potently inhibits *Ancylostoma ceylanicum* and *Necator americanus* hookworm larval development, with complete inhibition of development at 5 µg/mL (Figure 2A and B). The efficacy with Cry5B is very similar to that of Cry5B IBaCC (Figure 2A and B).

**Figure 2.**
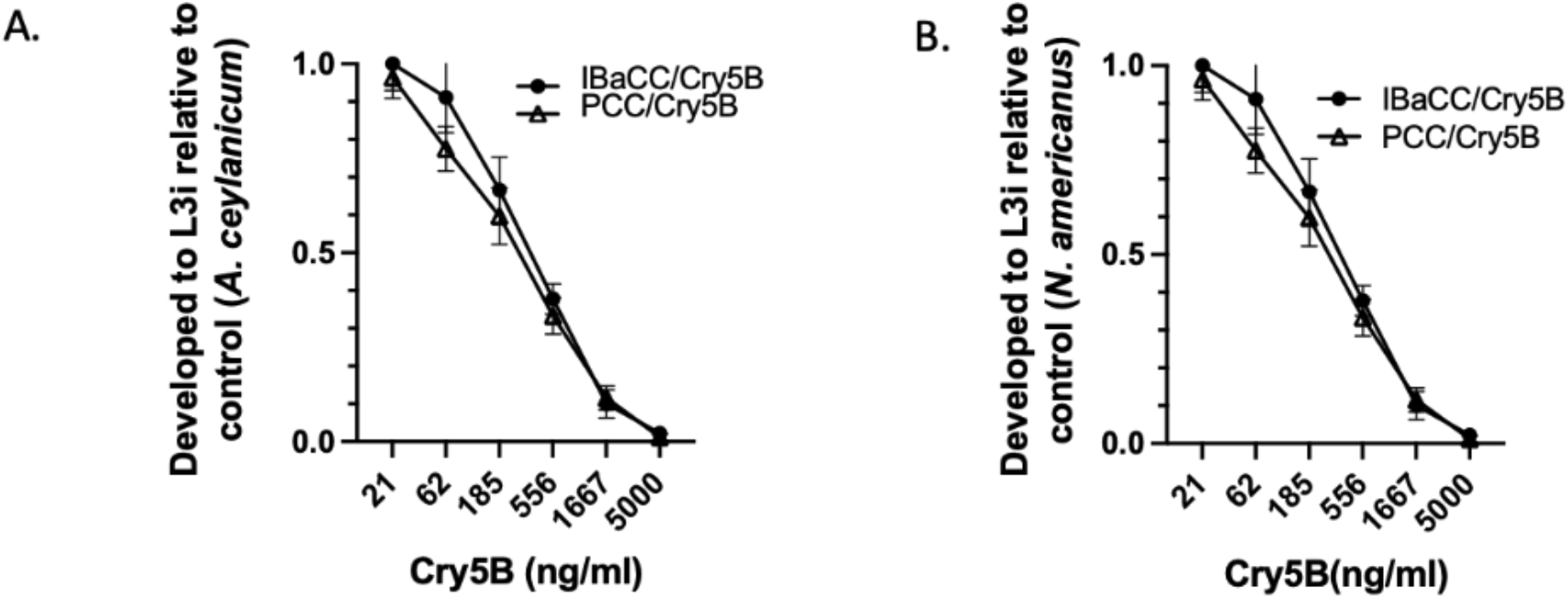
A comparison of the bioactivity of Cry5B IBaCC and PCC against developing hookworm larvae. (A) Ratio of *A. ceylanicum* hookworm eggs that developed to the L3i (infectious larval stage) in the presence of various doses of Cry5B presented as either IBaCC or PCC. (B) Ratio of *N. americanus* hookworm eggs that developed to the L3i (infectious larval stage) in the presence of various doses of Cry5B presented as either IBaCC or PCC. (A) and (B) represent the average of three independent trials. Error bars here and in all figures represent standard error of the mean.

We also tested Cry5B PCC against adult staged parasites *in vitro*. Cry5B PCC was found to be highly active *in vitro* against hookworm *A. ceylanicum* hookworm adults at two low concentrations. As a negative control for non-specific activity of purified crystals, we also expressed and purified insect-active Cry1Ac PCC, which had no detectable activity against these nematodes (Figure 3A). Good bioactivity of Cry5B PCC was also found against *N. americanus* hookworm adults (Figure 3B).

**Figure 3.**
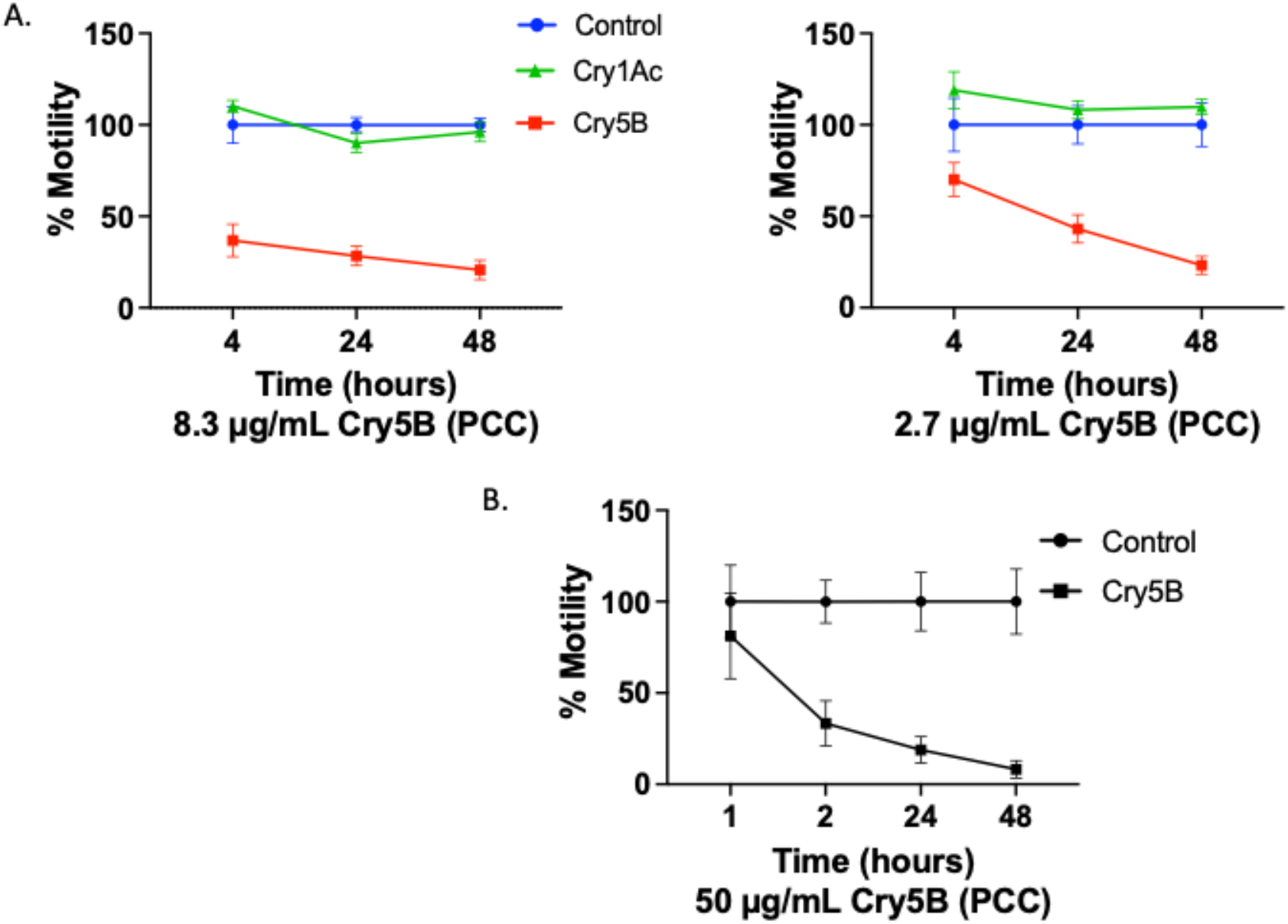
Intoxication of hookworm adults with PCC. (A) Adult *A. ceylanicum* hookworms were exposed *in vitro* to 8.3 µg/mL (left) or 2.7 µg/mL (right) Cry protein (Cry5B or Cry1Ac) as PCC. Motility was measured using the Worminator system (see Materials and Methods) and normalized to control (buffer only) at each time point. (B) Adult *N. americanus* hookworms were exposed to Cry5B PCC at 50 µg/mL and motility measured over time as for (A).

### Cry5B PCC is bioactive *in vivo* against an experimental hookworm infections

That Cry5B PCC was highly active against adult parasites *in vitro* suggested it might be efficacious against active parasitic infections *in vivo*. We therefore tested Cry5B PCC against an experimental hookworm infestation. Hamsters were experimentally infected with *A. ceylanicum* hookworms, a zoonotic parasitic hookworm that infects humans with re-emerging importance (30, 31). Eighteen days after infection, when the *A. ceylanicum* hookworms reached fertile adulthood, the hamsters were split into three groups based on equivalent fecal egg counts. Via oral gavage, one group received buffer control, one group received single dose of Cry5B IBaCC in buffer (final dose 5 mg/kg of Cry5B per hamster) and one group received a single dose of Cry5B PCC in buffer (final dose 5 mg/kg of Cry5B per hamster). Four days later, both parasite burdens and reproduction (fecal egg counts) were measured, as well as change in weight from the day of treatment until the end of the study. Cry5B PCC and IBaCC treatments respectively resulted in 74% and 82% *in vivo* reductions in parasitic hookworm burdens relative to buffer control (Figure 4A). The difference in efficacy based on parasite burdens between Cry5B IBaCC and PCC treatments was not statistically significant (P=0.55). Cry5B PCC and IBaCC treatments also respectively resulted in 63% and 77% reductions in parasite fecal egg counts (Figure 4B). The difference in efficacy based on parasite fecal egg counts between Cry5B IBaCC and PCC treatments was not statistically significant (P=0.33).

**Figure 4.**
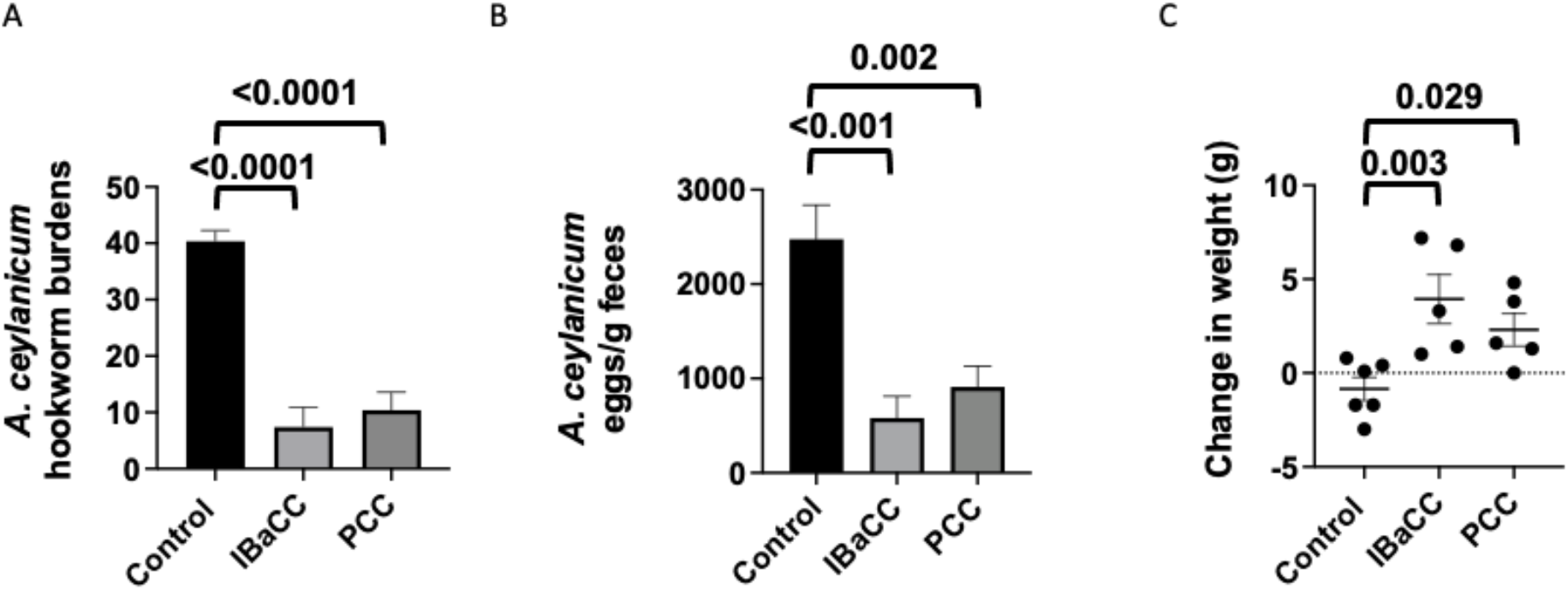
*In vivo* efficacy of Cry5B PCC against *A. ceylanicum* hookworm infections in hamsters. (A) Average intestinal hookworm burdens in hamsters treated with buffer, Cry5B as IBaCC (5 mg/kg), and Cry5B as PCC (5 mg/kg). Shown here and in other panels are P values for Cry5B treatment-group comparisons relative to control. n=6 for the control group and n=5 for each Cry5B-treated group. (B) Average fecal egg counts per gram of feces from the same hamsters as in (A). (C) Average change in weight from just before treatment until end of experiment from the same hamsters as in (A). Actual values are given in Table S1.

One of the sequelae of hookworm infection in humans is a negative impact on growth, which can also be seen in hamsters (32). We measured the change in weight in the four days between treatment and the end of the experiment (Figure 4C). Both treatment with Cry5B PCC and IBaCC on average resulted in hamsters that showed significant weight gain (+2.3 g and +3.9 g respectively) relative to buffer control, which showed weight loss (−0.85 g). The difference in weight gain between the PCC and IBaCC treated groups was not significant (P=0.33).

A small-scale *in vivo* efficacy trial of Cry5B PCC was also tested against *N. americanus* hookworm infections in hamsters. *N. americanus* is the dominant hookworm parasite of humans worldwide (27). Cry5B PCC was also highly effective against this genus of hookworms-- a single 10 mg/kg dose given orally to infected hamsters resulted in a 92% reduction in hookworm parasite burdens relative to buffer control and in a 98% reduction in parasite fecal egg counts (Figure 5A and B).

**Figure 5.**
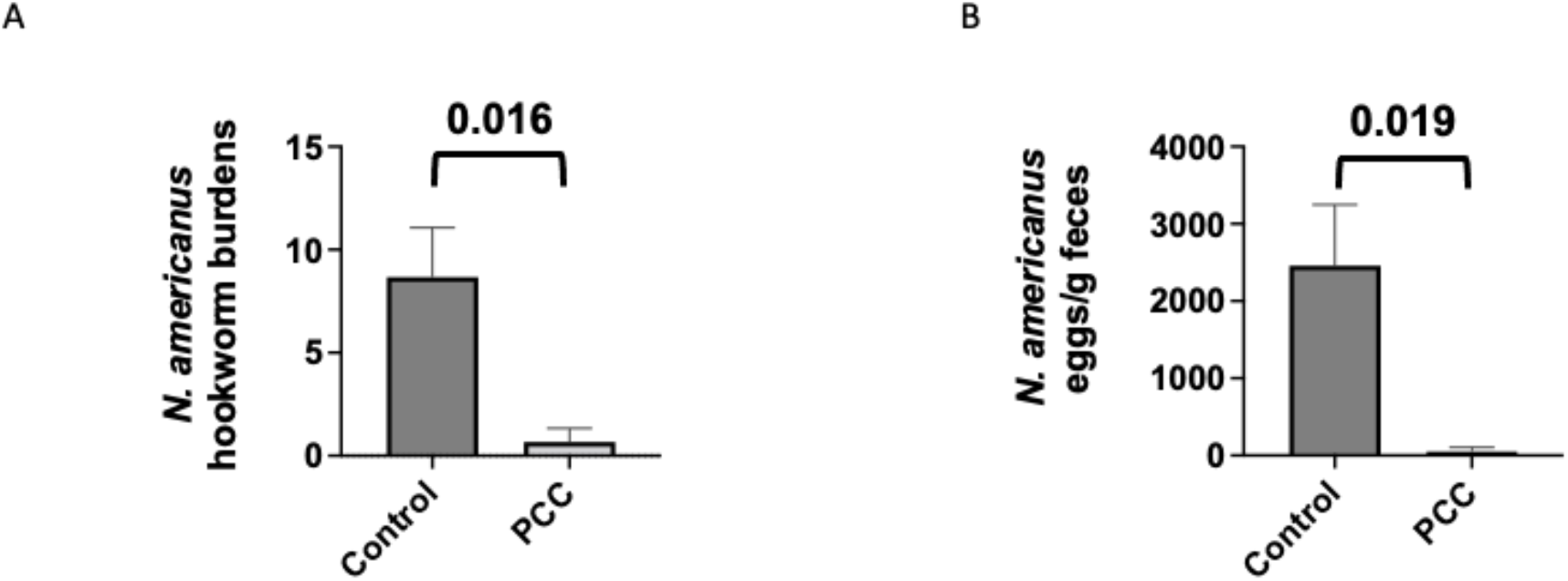
*In vivo* efficacy of Cry5B PCC against *N. americanus* hookworm infections in hamsters. (A) Average intestinal hookworm burdens in hamsters treated with buffer and Cry5B as PCC (10 mg/kg). n=3 for both groups. (B) Average fecal egg counts per gram of feces from the same hamsters as in (A). Actual values are given in Table S1.

### Cry5B PCC is effective against Ascarids including scale up and pilot study in foals

*Ascaris lumbricoides* or intestinal large roundworm is one of the most common parasites of humans worldwide and highly related to, if not the same species as, *Ascaris suum*, the dominant intestinal nematode parasite of pigs (33–35). We have reported on an immunodeficient mouse-model for *Ascaris suum* in which *A. suum* eggs gavaged into Th2-deficient mice develop into early intestinal parasitic L4 stage, which can then be used for intestinal *Ascaris* studies in rodents (17). Here, we tested Cry5B PCC against this experimental infection (Figure 6). A single 15 mg/kg dose of Cry5B PCC given via gavage led to a statistically significant 54% reduction in intestinal *A. suum* burdens in mice, similar to that of a single 15 mg/kg dose of Cry5B as IBaCC (57% reduction).

**Figure 6.**
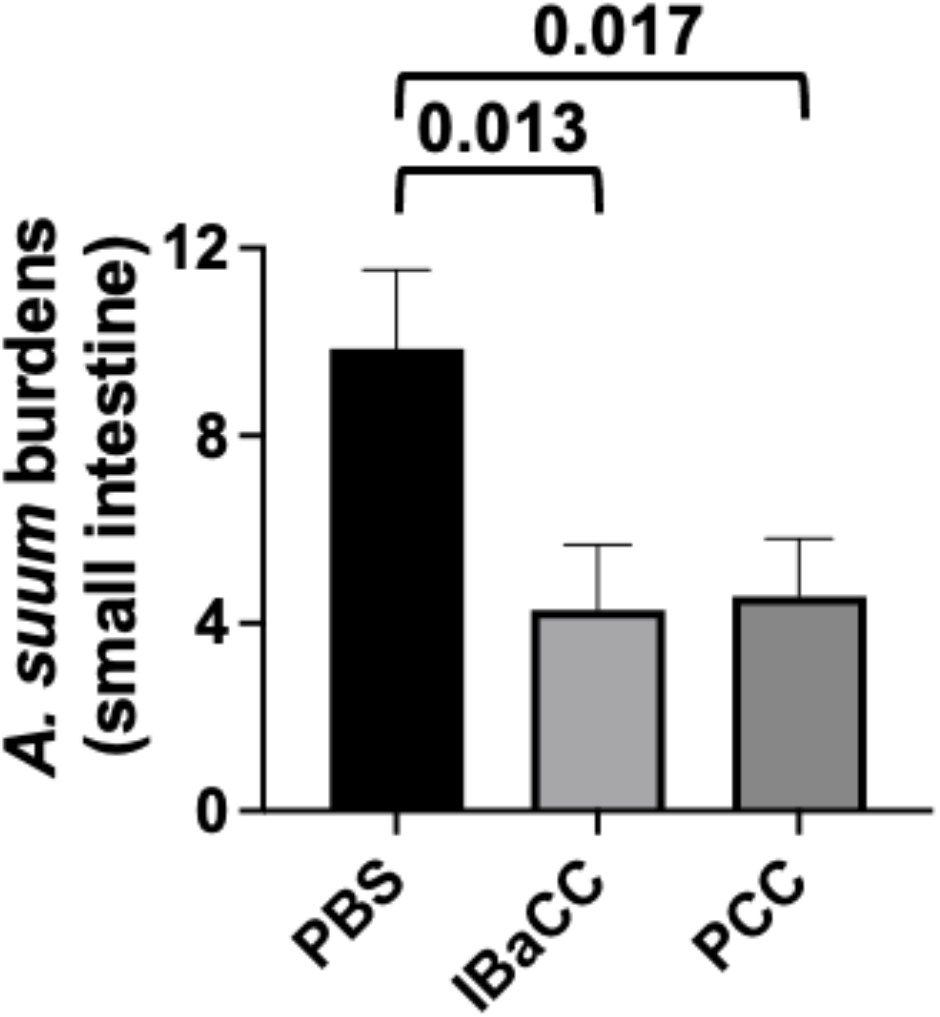
*In vivo* efficacy of Cry5B PCC against *A. suum* roundworm infections in mice.(A) Average intestinal *Ascaris* burdens in mice treated with buffer and Cry5B as IBaCC (15mg/kg) or as PCC (15 mg/kg). Shown here are P values for Cry5B treatment-group comparisons relative to control. n=7 for each group. Actual values are given in Table S1.

We have previously demonstrated that Cry5B is effective in foals against infestations of *Parascaris* spp., a critical and potentially lethal parasite of young horses related to human *Ascaris* (17, 36). We performed a pilot study in foals infected with *Parascaris* spp. to see if enough bioactive PCC could be produced for a large animal trial. Cry5B IBaCC was produced at 100 liter scale and then processed to PCC. Two foals, which had naturally acquired *Parascaris* spp. infections were enrolled in the study. After 6 weeks of monitoring, one foal was pretreated with omeprazole to neutralize stomach acid and then given a single oral 10 mg/kg dose of Cry5B PCC (2.5 g Cry5B as PCC). The second infected foal was kept untreated as a control. One week later (week 7), whereas the untreated foal retained its positive fecal egg count, for the PCC-treated foal, fecal egg counts dropped to zero and stayed at zero for the duration of weekly testing (weeks 8-11; Figure 7). The following week (week 12), the untreated control foal that had retained positive ascarid fecal egg counts was pretreated with omeprazole and then given a single oral 10 mg/kg dose of Cry5B PCC. One week later (week 13), fecal egg counts dropped to zero for this foal and stayed at zero for the duration of testing (week 14; Figure 7).

**Figure 7.**
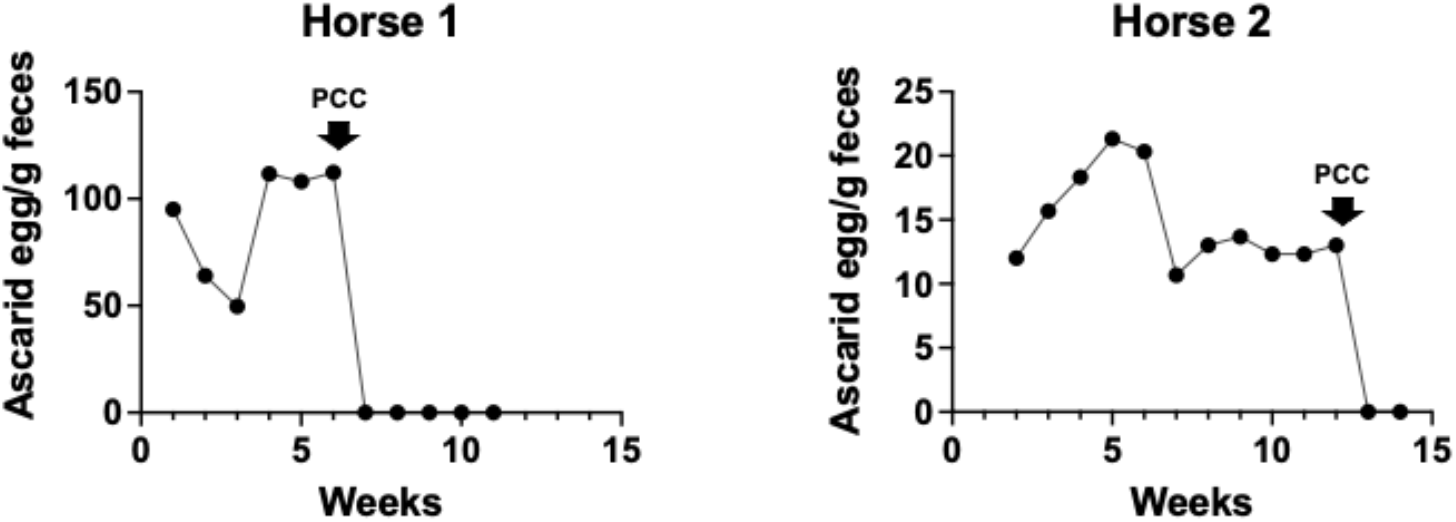
*Parascaris* fecal egg counts from two foals treated with scaled-up Cry5B PCC. Egg counts were taken weekly before and after treatment. Treatment occurred immediately after fecal samples were taken on the week indicated by the down arrow.

## DISCUSSION

Here we describe a scalable method for purification of crystals from sporulation-defective *Bacillus thuringiensis*. In a strain expressing Cry5B, protein content in these Purified Cytosolic Crystals (PCC) was >95% Cry5B. Cry5B PCC had excellent bioactivity as indicated by its potent *in vitro* effects on larval and adult stages of two species of human hookworms. Cry5B PCC is also highly effective at clearing two different species of human hookworm infections in infected rodents and two different species of ascarids, one in rodents and one in horses. As part of the equine study, scale up to 100 liters and purification of gram quantities of Cry5B PCC was achieved. These data indicate PCC is scalable, highly active, and a potent anthelmintic.

IBaCC, from which PCC is derived, introduced the concept of using a dead bacterium to deliver live Bt Cry proteins. PCC takes that process one-step further. In instances where increased purification of Bt Cry proteins is desired (see below), PCC offers significant improvements over other technologies. In addition, in the case of anthelmintic treatment of humans and animals, PCC also offers an active pharmaceutical ingredient (API) highly compatible with drug delivery, eliminating the bacterial shell surrounding IBaCC and greatly increasing the purity of Cry5B protein. We were able to also produce PCC from a strain expressing insecticidal Cry1Ac. The PCC process can thus be extended to a wide range of Bt three-domain proteins. Studies on bioactivity of Cry1Ac PCC against insects are in progress (MM and DL, manuscript in preparation).

Purified Cry proteins have a number of potentially important applications (37). They can be used for crop protection against target insects devoid of live bacteria and bacterial products. This is becoming increasingly important as there are indications that live Bt spore crystal lysates used as topical sprays might be associated with adverse health events and are subject to increasing regulation (see Introduction). Given its purity, PCC is also ideal for making formulations or concentrating Cry protein products to increase specific activity. PCC could be used for producing antibodies for immunoassays for detection of transgenic plants. PCC could also be used in insect and nematode bioassays to determine the effects of a purified Cry protein independent of other potential protein toxins produced by Bt. Importantly, PCC could also be useful for biosafety studies required for use of Cry proteins in transgenic plants and as anti-parasitics. The additional step of DNAse, which results in increased solubility of the crystals, can be useful for using PCC to study the effects of pure Cry proteins against targets for which uptake of protein potentially requires dissolution of the crystals, *e*.*g*., nematodes that restrict intake of large proteins/particles such as some plant-parasitic nematodes and whipworms (38, 39).

The use of hexane here to process the lysed IBaCC and achieve phase-partitioning to aid in crystal purification has also been reported for processing spore-crystal lysates and removal of spores (40). The process described here has several advantages over the previous work. First, the PCC process achieves many orders of magnitude greater reduction in colony forming units/mL. Second, the PCC process results in a much greater percent Cry protein purity in protein content.

In summary, here we demonstrate a scalable and simplified procedure to produce a highly purified form of Bt crystals called PCC for use in a diverse range of applications (*e*.*g*., drug development, concentration of bioactivity, diagnostics, safety studies, bioactivity studies, formulation for environmental release) where use of purified Bt crystals, soluble Cry proteins are desirable or preferred.

## MATERIALS AND METHODS

### Preparation of PCC

BaCC (**B**acteria with **C**ytosolic **C**rystal(s)) expressing either Cry5B or Cry1Ac were grown and treated with essential oil as previously described to make IBaCC (14). The bacterial cells were lysed using 200 ug/ml Lysozyme in buffer (20 mM Tris-HCl, pH 8.0, 2 mM EDTA, 1 % Triton X-100) for 2-4 h at 37 °C. After the lysozyme treatment the cells were subjected to sonication on ice until 95% bacterial lysis was observed using a UPlanFl 100x/1.30 Oil Ph3 objective on an Olympus BX60 microscope. Soluble bacterial proteins and cellular constituents were removed from the Cy5B crystals by centrifugation at 4500 rpm, 2 h, 4 °C in Eppendorf benchtop centrifuge 5804R as the dense crystals are pelletable. Since DNA is known to stabilize crystals (see above), an optional DNase treatment can be performed based on manufacturer’s instructions on the resuspended pellet to increase the solubility of Cry5B in PCC *in vitro* with no impact on bioactivity. The crystal containing pellet was resuspended in water and then brought to 1 M NaCl. Hexane or food grade oil (*e*.*g*., corn oil) was added to a final concentration of 25% and incubated for 2 h at room temperature in a rotating rotor after pulse sonication to ensure disruption of crystals from cell debris. Following centrifugation, the crystal containing pellet fraction was washed thrice with sterile water to remove hexane or oil and finally resuspended in sterile water (one-tenth volume). These stable PCC aliquots were stored in -80 °C and subjected to various QC analysis and *in vitro* and *in vivo* worm toxicity assays. Microscopy of IBaCC and PCC was carried out using a 100x bright field microscope. To determine the percent Cry5B protein in IBaCC and PCC, Cry5B IBaCC and PCC were subjected to sodium dodecyl-sulfate polyacrylamide gel electrophoresis (SDS-PAGE), stained with Coomassie Blue, and destained. The gels were imaged using the BIORAD ChemiDoc™ XRS+ Molecular Imager^®^ Imaging System. The intensity of the full-length Cry5B was then compared to the intensity of the protein bands below the full-length Cry5B band. This was done by comparing the integrated density value of the PCC lane and the IBaCC lane. This value was calculated using the most current ImageJ software. This calculation was repeated at various concentrations of IBaCC and PCC, with very similar results obtained at each.

### Animals and parasites

*Ancylostoma ceylanicum* and *Necator americanus* hookworm life cycles were maintained in hamsters as previously described (15) with the exception that *N. americanus* was maintained with use of 4 mg/L dexamethasone in the drinking water without any need for supplemental dexamethasone injection. Hamsters were provided with food and water (ad libitum). All animal experiments involving hookworms were carried out under protocols approved by the University of Massachusetts Chan Medical School (PROTO202000071/A-2483). All housing and care of laboratory animals used in this study conform to the NIH Guide for the Care and Use of Laboratory Animals in Research (18-F22) and all requirements and all regulations issued by the USDA, including regulations implementing the Animal Welfare Act (P.L. 89-544) as amended (18-F23). All protocols were approved by the University of Kentucky Institutional Animal Care and Use Committee (Protocol 2015–2078) for the equine studies and the USDA Beltsville - Institutional Animal Care and Use Committee #17–019 for the murine studies along with Institutional Biosafety Committee #271.

Egg-to-L3i (infectious third larval stage) assays (Figure 2) were carried out as described (29). For both Figure 2A and B, the data were normalized to the number of eggs out of ∼60 in each well that developed to the L3i stage in the absence of any Cry5B (IBa (14) as the control for IBaCC and water as the control for PCC). For Figures 2A and 2B, 100% represents between 23-27 eggs that developed to L3i, depending upon the negative control and parasite.

Adult hookworms harvested from the intestines of infected hamsters were assayed *in vitro* as previously reported (14) with the exception that motility was read out using the Worminator platform (41), averaging over 3 minutes with one adult parasite per well. For *A. ceylanicum*, the data represent the average of two independent experiments with n=8 adults per dose per experiment. For N. americanus, 8 adults were used in the experimental group and 7 were used in the control. For each *in vitro* experiment, motility was normalized to that of the buffer control at each time point. For *A. ceylanicum* 8.7 ug/mL, control values were 74.4, 91.9 and 78.5 at 4, 24 and 48 hours respectively. For *A. ceylanicum* 2.3 ug/mL, control values were 58.5, 82.8 and 74.1 at 4, 24 and 48 hours respectively. For *N. Americanus*, control values were 78.0, 68.2, 87.2, 85.3 at 1, 2, 24, and 48 hours respectively.

*In vivo* hookworm curative assays were carried out essentially as described (14). For *A. ceylanicum*, hamsters were orally infected with 140 infectious larvae. An overnight collection day 18 PI was taken to determine pre-treatment fecal egg counts (FECs). These FECs were used to place hamsters into similarly infected groups. Following grouping, hamsters were weighed and then orally treated with either 0.1 M bicarbonate buffer, 5 mg/kg Cry5B IBaCC mixed with 200ul of 0.1 M bicarbonate buffer, or 5mg/kg Cry5B PCC mixed with 200ul of 0.1 M bicarbonate buffer. Bicarbonate buffer is used to partly neutralize stomach acid and itself has no effect on hookworm burdens (14). Day 22 PI, an overnight collection was taken; day 23 PI, animals were weighed, and then euthanized. Total intestinal hookworm burdens and FECs were determined as was the change in weight. For *N. americanus*, hamsters were infected subcutaneously with 300 infectious larvae; the curative experiment was otherwise carried out as described (14).

Curative studies of *A. suum* infections in mice were carried out similar to as already described (17), except IL13-knockout mice in a C57BL/6 background were used instead of STAT6-knockout mice (The IL13-KO mice were supplied as a gift from the laboratory of Thomas Wynn through an NIH contract with Taconic). Male and female mice of 3-4 months of age were orally inoculated with 1500 *A. suum* eggs. Twelve days PI, mice were orally gavaged with either phosphate buffered saline (control), IBaCC (Cry5B 15 mg/kg), or PCC (Cry5B 15 mg/kg), all as a single dose. Sixteen days PI, mice were euthanized and L4 *A. suum* burdens in the small intestine were assessed.

A pilot curative study in foals was carried out as follows. Two foals naturally infected with *Parascaris* spp. were identified based on fecal egg counts determined by the Mini-FLOTAC technique (42). The timing of weekly fecal egg counts is provided in the figure. One foal was initially treated and the other kept as a control. To maximize the efficacy, the first foal to be treated was pretreated with 4.0 mg/kg omeprazole (Gastrogard, Boehringer Ingelheim, Ingelheim, Germany) administered orally to neutralize stomach acid and allow more Cry5B to pass into the small intestine. Sixteen hours post-treatment with omeprazole, the foal received PCC via nasogastric tube at a Cry5B dose of 10 mg/kg (∼2.5 g/horse). Two weeks after the first foal was removed from the trial, the second foal, which had retained fecal egg counts, was similarly treated with omeprazole and Cry5B PCC.

### Statistical Methods

All statistical analyses and graphs were generated using GraphPad Prism Version 9.3.1. All comparisons involving three groups in which two groups were each compared to a buffer control were analyzed using One-Way Analysis of Variance (ANOVA) using one-tailed Dunnett’s post-test to test the hypothesis that treatment would result in a lower worm burden, lower fecal egg count, or increased weight relative to control. All comparisons between two groups were carried out using two-tailed Student’s t-test (non-parametric tests such as Mann-Whitney could not be used because of sample size limits, *e*.*g*., three hamsters per group in the *N. americanus* curative experiment).

## ACKNOWLEDGEMENTS

This work was financially supported by (1) the National Institutes of Health/ National Institute of Allergy and Infectious Diseases grants R01-AI056189 and R01-AI150866 to R.V.A., (2) Agriculture and Food Research Initiative Competitive Grant no. 2015–11323 from the USDA National Institute of Food and Agriculture to R.V.A.

---

**Table S1.**
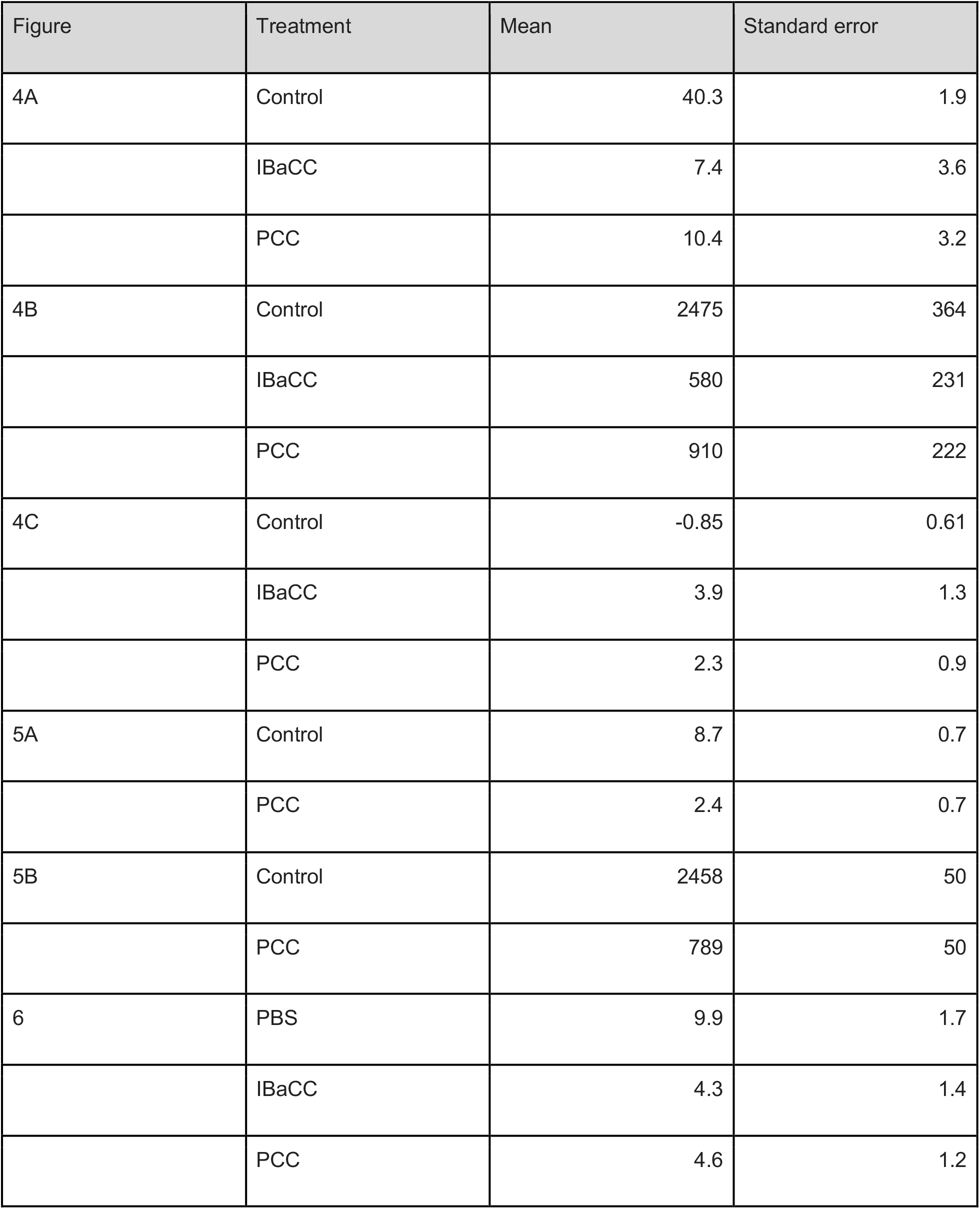
Data from rodent *in vivo* experiments

